# Unpacking similarity effects in visual memory search: categorical, semantic, and visual contributions

**DOI:** 10.1101/2025.04.09.647918

**Authors:** Linlin Shang, Lu-Chun Yeh, Yuanfang Zhao, Marius V. Peelen

## Abstract

Visual memory search involves comparing a probe item against multiple memorized items. Previous work has shown that distractor probes from a different object category than the objects in the memory set are rejected more quickly than distractor probes from the same category. Because objects belonging to the same superordinate category usually share both visual and semantic features compared with objects of different categories, it is unclear whether the category effects reported in previous studies reflected visual and/or semantic target-distractor similarity. Here, we employed old/new recognition tasks to examine the role of categorical, semantic, and visual similarity in short- and long-term memory search. Participants (N=64) performed visual long-term memory (LTM) or short-term memory (STM) search tasks involving animate and inanimate objects. Trial-wise RT variability to distractor probes in LTM and STM search was modelled using regression analyses that included predictors capturing categorical target-distractor similarity (same or different category), semantic target-distractor similarity (from a distributional semantic model), and visual target-distractor similarity (from a deep neural network). We found that categorical, semantic, and visual similarity all explained unique variance in trial-wise memory search performance. However, their respective contributions varied with memory set size and task, with STM performance being relatively more strongly influenced by visual and categorical similarity and LTM performance being relatively more strongly influenced by semantic similarity. These results clarify the nature of the representations used in memory search and reveal both similarities and differences between search in STM and LTM.

## Introduction

When going to a supermarket to buy beverages for dinner, you need to compare the beverage in front of you with your shopping list in long-term memory (LTM). This example of probe-recognition memory illustrates how we frequently need to recognize whether an object in view (the probe) is one we memorized previously. Interestingly, previous work has shown that visual memory search is easier for some probe objects than others. For example, when the memory set consists of objects that all belong to one category (e.g., milk, beer, fruit juice – all beverages), participants are slow to decide that an object from the same category (e.g., cola) is not on the memory list relative to an object from another category (e.g., a plant; Cunningham & Wolfe, 2012, 2014; Drew & Wolfe, 2014; Gronau et al., 2024; Shang et al., 2024; Williams et al., 2024). In these studies, participants first memorized a varying number of visually presented objects that all belonged to one category (e.g., animals) and then performed an old/new probe recognition task in which they decided whether visually presented objects were part of the memory set (i.e., targets) or not (i.e., distractors). The studies found that participants were much faster to reject distractor probes coming from a different (non-target) category (e.g., inanimate objects) than they were to reject distractor probes coming from the same (target) category, particularly when memory set size was high. In the current study, we investigated what drives this category effect, considering three factors that could contribute to it: visual similarity, semantic similarity, and target category-level selection.

First, the category effect could reflect differences in the visual similarity between items within versus between categories: a milk bottle and a cola bottle are visually more similar than a milk bottle and a plant. Visual similarity is known to strongly influence attentional selection in visual search tasks (Duncan & Humphreys, 1989). Considering the close relationship between attention and memory (for a review, see Sherman & Turk-Browne, 2024), visual similarity could also influence memory search, such that visually distinct items are more quickly rejected during memory search. Evidence that LTM contains information about visual object features comes from studies showing that participants can accurately memorize precise details of large numbers of images differing only in subtle visual or state/pose details (e.g., Brady et al., 2008; Vogt & Magnussen, 2007). For example, Vogt and Magnussen (2007) had participants view 400 photographs of doors, and found that participants were able to accurately discriminate between previously seen and unseen doors, even up to a week later, demonstrating that visual LTM has a high capacity and stores detailed visual information. Accordingly, the relatively slow rejection of same-category distractors in memory search may reflect the visual similarity of these distractors to the items in the memory set.

Second, the category effect could reflect differences in the semantic (or conceptual) similarity between items within versus between categories: a milk bottle and a cola bottle are semantically more closely related than a milk bottle and a plant. In line with a role for semantic similarity, memory search is often compared to a dynamic “foraging-like” search (Davelaar, 2015; Hills et al., 2012, 2015; Hills & Pachur, 2012), such that items related to the probe stimulus are concurrently activated in memory based on semantic similarity to generate search patches and facilitate memory search (Collins & Loftus, 1975; Collins & Quillian, 1969). There is evidence that even visual LTM may depend more on conceptual than perceptual similarity (Konkle et al., 2010a, 2010b). For example, in a study that tested visual LTM for objects, memory performance for an object was worse when the memory set had included multiple exemplars of that object’s category. Importantly, however, this effect was better explained by the conceptual than the perceptual similarity of the same-category objects in the memory set (Konkle et al., 2010b). Similarly, the previously observed benefit of different-category distractors in memory search may primarily reflect the semantic distinctiveness of these distractors relative to the items in the memory set.

Finally, the category effect could reflect a binary task-defined categorical effect. Specifically, when all items in the memory set come from one category, participants may use that information to select which objects should enter memory search. For example, while searching for the three beverages, participants may first decide whether an object belongs to the target category – whether it is a beverage or not – before commencing memory search. It should be noted that deciding whether an object is from the target category or not may itself be influenced by the object’s visual and semantic similarity to the target category, but categorical similarity in this case is binary (target vs non-target category) and therefore not identical to visual and/or semantic similarity, which are continuous (e.g., objects within a category vary in terms of their visual and semantical similarity). Furthermore, target category selection is flexible and task-dependent and can, in principle, exist independently of general visual and semantic similarity. For example, when searching for the three beverages, a participant may categorically reject all beverages that are outside of their budget. The first and primary aim of the current study was to examine the distinct contributions of visual and semantic similarity, as well as categorical selection, to memory search performance.

Our second aim was to compare these contributions across visual LTM and visual working memory/short-term memory (WM/STM). Visual STM (VSTM) is a visual storage that supports the maintenance of visual information over a brief interval (up to a couple of seconds). The capacity of VSTM is much smaller than the capacity of LTM and the format of VSTM is likely more closely related to visual perception than the format of LTM (for reviews, see Luck, 2008; Schurgin, 2018) Evidence for the visual nature of VSTM comes from neuroimaging studies, showing that VSTM relies on activity in visual cortex. For example, Harrison and Tong (2009) found that activity patterns in early visual cortex during the memory delay predicted the orientation of visual gratings held in STM. More generally, visual cortex representations may be activated (“held online”) during visual STM but not LTM (see Sreenivasan et al., 2014 for a review). This suggests that STM representations are stored in a more visual code than LTM representations and predict that STM may be more strongly influenced by visual similarity than LTM. Conversely, LTM may be more strongly influenced by semantic similarity than STM.

To investigate distinct contributions to memory search performance, participants (N=64) performed visual long-term memory (LTM) or short-term memory (STM) search tasks involving animate and inanimate objects. To measure the category effect, the items in the memory set always came from one category (animate or inanimate) while the probe objects came from both categories. Because probe objects from a different category than the memory set can only be distractor objects (i.e., objects not in the memory set), our main analyses focused on reaction times (RTs) to distractor objects. This ensured that the response (“no”) was matched across target and non-target categories. Specifically, we modelled trial-wise RT variability to distractor probes using regression analyses that included predictors capturing categorical target-distractor similarity (same or different category), semantic target-distractor similarity (from a distributional semantic model), and visual target-distractor similarity (from a deep neural network). This approach allowed us to investigate: 1) whether the category effects reported in previous studies remain when regressing out visual and semantic similarity, 2) whether visual and semantic similarity independently contribute to memory search performance, and 3) whether the contributions of categorical selection, and semantic and visual similarity to memory search performance differ between LTM and STM search.

## Method

### Data availability

Analysis code and stimuli are available at https://github.com/shangll/OData_VisSimi; data at: https://osf.io/9zbq4/

### Participants

In total, sixty-four participants were recruited from the online platform Prolific and signed an online informed consent form. Thirty-one participants (18 male and 13 female) between 20 and 35 years of age (*M* = 23.80, *SD* = 2.92) participated in the LTM task while the other thirty-three new participants (25 male and 8 female) between 18 and 35 years old (*M* = 23.76, *SD* = 3.08) were recruited for the STM task. In total, one participant in the LTM task and three participants in the STM task were excluded because of low accuracy in the set size 8 conditions (below 3 *SD* of the group mean), preventing accurate estimation of reaction times (our main dependent measure). All participants received 6 euro per hour after the experiment. The study was approved by the Ethics Committee of the Faculty of Social Sciences, Radboud University Nijmegen.

### Stimuli and design

A total of 1020 full-colour images including 30 animate categories and 30 inanimate object categories were selected from Google Images and from Brady et al. (2008), respectively. Each category contained 17 exemplar images. Each image was used only once in each task. The current study followed a 2 (LTM/STM task; between-subjects) × 2 (within/between category; within-subjects) × 4 (memory set size (MSS) 1/2/4/8; within-subjects) mixed design. For each task, all participants were tested on both categories over all four MSS. Both tasks were programmed with PsychoPy v2020.2.3 (Peirce et al., 2019) and were hosted on Pavlovia. Before each task, 20 practice trials were conducted to ensure that the participants were familiar with the experimental instructions and procedure.

### Procedure

#### LTM task

The LTM task was an old/new recognition task with 50% target-absent trials. The experiment was structured in blocks in which one MSS was used. Each block consisted of a study phase, a memory test phase, and the main experiment phase. In the study phase, a central fixation cross appeared for 0.5 sec as a prompt indicating the beginning of the block, followed by 1, 2, 4, or 8 targets from either animate or inanimate category presented in the centre of the screen on a white background, one at a time, for 3 sec with an inter-stimulus interval of 0.95 sec (Figure 1A). Participants were instructed to memorize the objects. The subsequent memory test phase followed the same procedure except for the duration of the object display. In addition, when the test object was presented, participants needed to decide whether it was part of the memory set (target) or not, and the display terminated upon response. Only participants with >80% accuracy twice consecutively were allowed to continue to the next and main phase.

**Figure 1.**
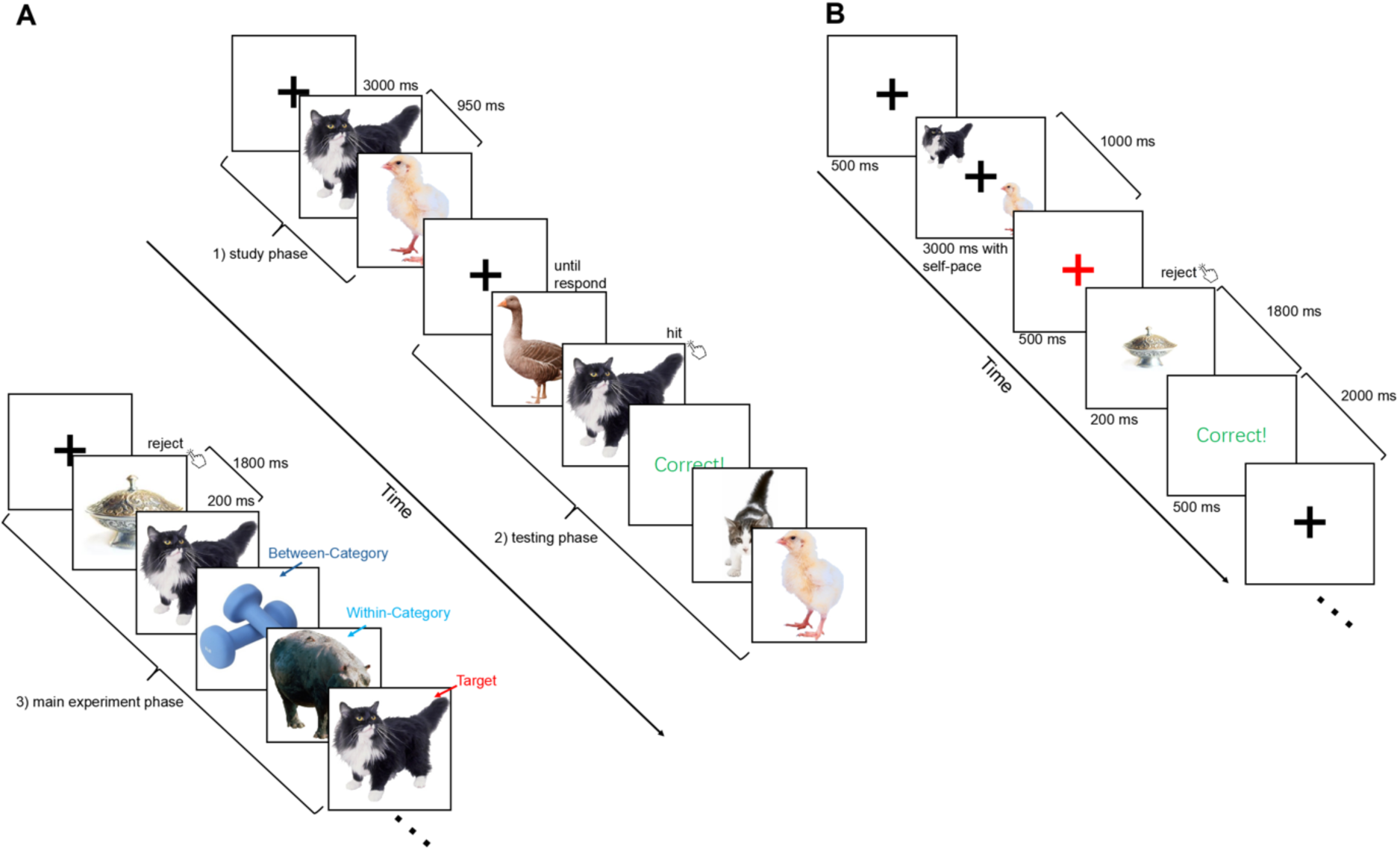
Illustrations of the procedures of (A) LTM and (B) STM Tasks. *Note*. Examples of an animate condition with two target objects (i.e., MSS 2). (A) In a study phase, participants in the LTM task first memorized two objects from the same category (the animal category in this example). Once they memorized the objects, as verified in the testing phase, they continued with the main experiment (lower left panel), deciding if a probe object was a target or not (old/new recognition task). Distractor objects could come from the same category (within-category distractor) or from the other category (between-category distractor). All analyses focused on reaction time to the distractor objects. (B) As in the LTM task, participants in the STM task also memorized one or multiple objects (here: two), but these were presented simultaneously and were directly followed, after a 1.5-sec retention interval, by a single probe object. A new memory set was presented on the next trial.

The main experiment phase consisted of eight blocks of 60 trials each (30 targets and 30 distractors (15 animate and 15 inanimate). A central fixation cross appeared for 0.5 sec, followed by a target or a distractor object presented in the centre of the screen on a white background for 0.2 sec and then an inter-trial fixation interval of 1.8 sec. Participants were required to decide whether the object presented on the screen was a target or not within 2 sec. All distractors were only shown once in the experiment. All targets were only used in one block, but the targets could repeat within a block. Trial order was randomized.

#### STM task

For the visual STM task, the stimuli were randomly selected from the same stimulus pool as used in the LTM task, and were equally divided into 7 blocks. Each block consisted of 32 trials (16 target and 16 distractor probes (8 animate and 8 inanimate)). Participants could take a brief break at the end of each block.

A black central fixation cross appeared for 0.5 sec as a prompt indicating the beginning of the trial, and then participants were required to memorize 1 to 8 objects from either animate or inanimate category. These objects were presented simultaneously for a maximum of 2, 3, 5, or 9 sec depending on the memory set size (1, 2, 4, 8 items, respectively); participants could shorten this by pressing a button. After a 1-sec fixation interval, the fixation cross turned red for 0.5 sec and a single probe object was presented for 0.2 sec, followed by a 1.8-sec response window (or until response). Participants had to respond within 2 sec or they would get negative feedback. The inter-trial interval was also 2 sec (Figure 1B).

#### Behavioural data analyses

For both tasks, the main analyses focused on RT of target-absent (i.e., distractor) trials. Only correct trials were included in RT analyses. Trials on which participants failed to respond were counted as incorrect. For each set size and each category condition, RTs beyond 3 standard deviations (SD) of the mean were excluded as outliers. Under this criterion, 1.364% of data points in the LTM task and 1.136% of data points in the STM task were excluded from further analyses on RT.

#### Overview of similarity-based analyses

To understand how visual, semantic, and categorical similarity contribute to memory performance, a similarity-based analysis was employed. The key idea was to quantify the relationships between pairs of visual objects in terms of their visual features, semantic meaning, and task-related category (animate/inanimate). We then investigated how these quantified similarities predict behavioural performance (i.e., RTs) in both short-term (STM) and long-term memory (LTM) tasks. First, neural networks were used to establish visual and semantic similarity (see below), while a binary measure was used to indicate whether two images belonged to the same or different category (e.g., animate vs inanimate). Then, generalized linear models (GLM) were employed to predict RTs with each type of similarity (visual, semantic, and categorical). Finally, stepwise regression (i.e., semi-partial) was employed to disentangle the unique contribution of each variable. We performed these analyses separately for our LTM and STM tasks, enabling comparisons of how visual, semantic, and categorical similarities differentially impact memory search performance.

#### Visual similarity

AlexNet (Krizhevsky et al., 2017) is a commonly used neural network model of human visual object processing, showing good alignment with visual cortex representations (Jozwik et al., 2018). We used AlexNet to establish the visual similarity between all pairs of objects.

Specifically, we used Spearman correlation to correlate the flattened activation matrices of each pair of objects at each layer, and thereby obtained eight representational similarity matrices (one for each layer). In Layer 6, the pair with the highest visual similarity (excluding pairs within the same subcategory) was fish and bowtie (ρ = 0.610) while the pair with the lowest visual similarity (excluding pairs within the same subcategory) was abacus and kangaroo (ρ = -0.156).

Next, we used these correlations to establish the visual target-distractor similarity at the single trial level, in order to relate this measure to behavioural performance across trials. In both tasks, however, in a given target-absent trial, the individually presented distractor object could be paired with multiple targets (when MSS>1). Thus, for each target-absent trial, we would obtain at least two correlation coefficients for each layer. Here, two approaches were employed to compute the visual similarity for each trial. The first approach was to average the similarity correlations of the multiple targets and the single distractor for each trial at each layer. The second approach was to select the maximum similarity correlation on a given trial, based on the idea that behavioural performance may be limited by the most similar item. However, we did not observe any statistical difference between these two approaches. Therefore, all reported results were based on the mean similarity correlation (see Supplemental Materials for the results of maximum similarity).

#### Semantic similarity

To compute semantic similarity, all object images were first given names at the basic category level (e.g., “cat”, “monkey”, “guitar”, “bag”). Three authors independently named the objects. If there was discrepancy, they discussed the names to come to a consensus. The semantic similarity of these names was then determined using Word2Vec (Mikolov et al., 2013; Řehůřek & Sojka, 2010), a method of natural language processing (NLP). Through modelling text within large corpora, Word2Vec uses word co-occurrence patterns to identify words with similar contexts and then maps them to vectors that can be considered in terms of their cosine similarity. The pair with the highest semantic similarity was shrimp and fish (ρ = 0.610) while the pair with the lowest semantic similarity was hippopotamus and telephone (ρ = -0.225). Trial-wise semantic similarity was computed in the same way as described above for visual similarity. Also here, mean and max similarity gave highly similar results and only mean similarity results are reported (see Supplemental Materials for the results of maximum similarity).

#### Mean similarity for within- and between-category pairs

Intuitively, objects of the same category are visually and semantically relatively similar. To test whether this was the case for the objects in our experiments, we employed AlexNet and Word2Vec to compute the visual similarity and semantic similarity between 1020 objects, both for within-category (e.g., frog-rabbit; both animate) and between-category (e.g., frog-sofa; animate-inanimate) pairs. Duplicate and auto-correlated pairs were excluded. Because semantic similarity was reflected at the subcategory level rather than at the individual image level (e.g., although the individual objects within the subcategory of ceiling fan showed distinct visual differences, they were all semantically labelled as “ceiling fan”), for this analysis, we averaged the similarity values for each subcategory (there were a total of 60 subcategories: 30 animate + 30 inanimate). This way, 1770 subcategory pairs were included (i.e., the number of unique elements in a symmetric 60 × 60 similarity matrix). The results (Figure 2) showed that from the 2^nd^ layer, visual similarity was significantly higher for within-than between-category pairs for each layer (see Table 1 for statistical values). The mean semantic similarities for within- and between-category pairs also differed: 0.173 and 0.066, respectively (permutation *t* = 19.932, *p* < 0.001). Finally, visual and semantic similarity were moderately correlated across subcategories (this correlation ranged from 0.068 in Layer 1 to 0.432 in Layer 6). Importantly, however, visual and semantic similarity were not identical and could be dissociated. For example, Layer-6 visual similarity between “frog” and “rabbit” was low (ρ = 0.07), while their semantic similarity was relatively high (ρ = 0.40).

**Figure 2.**
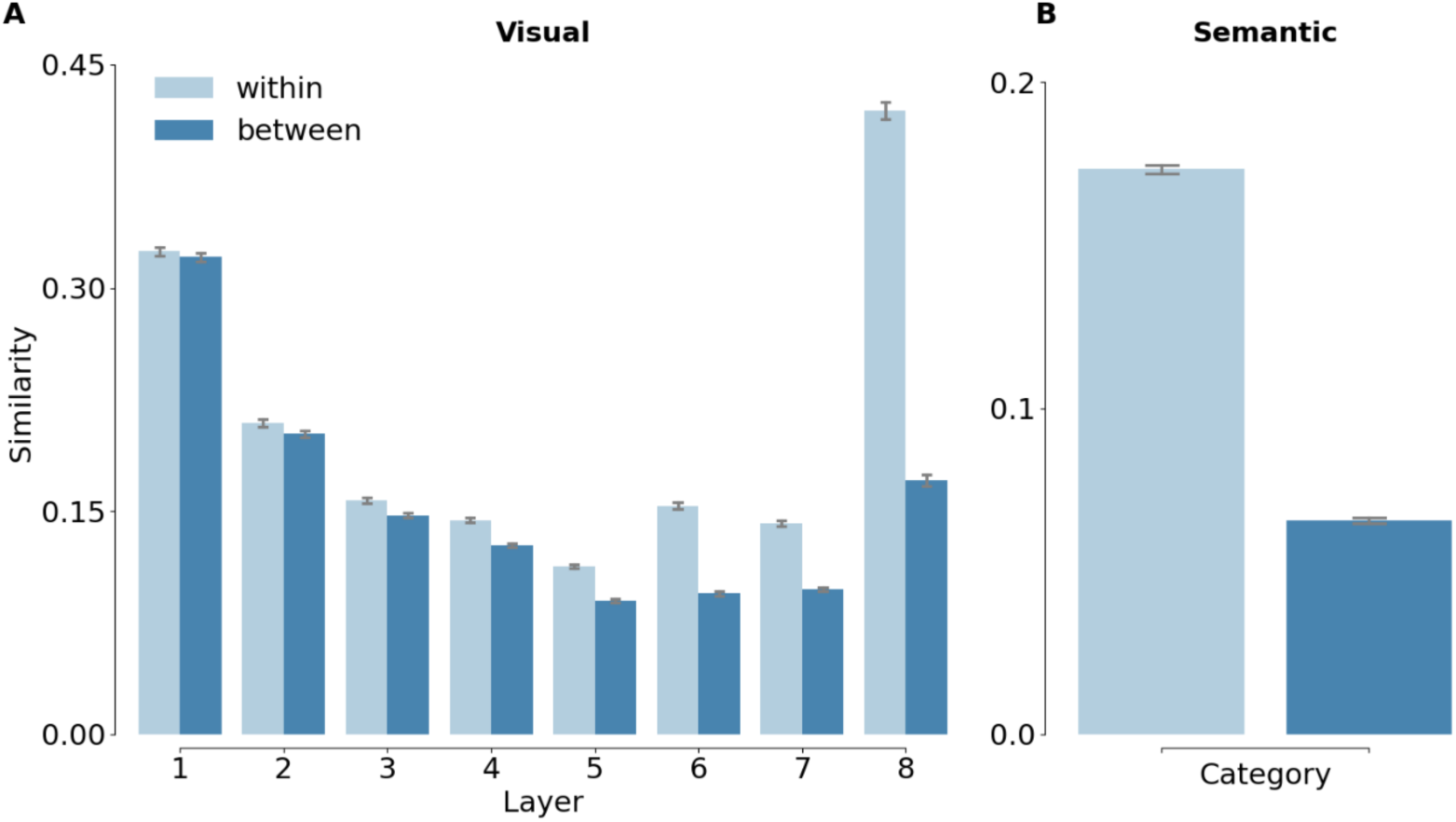
The differences in visual/semantic similarity between object images within the same category and across different categories. *Note*. For all figures, light blue bars indicate the within-category condition while the dark blue indicate the between-category condition. The error bars represent the standard error of the mean.

**Table 1.**
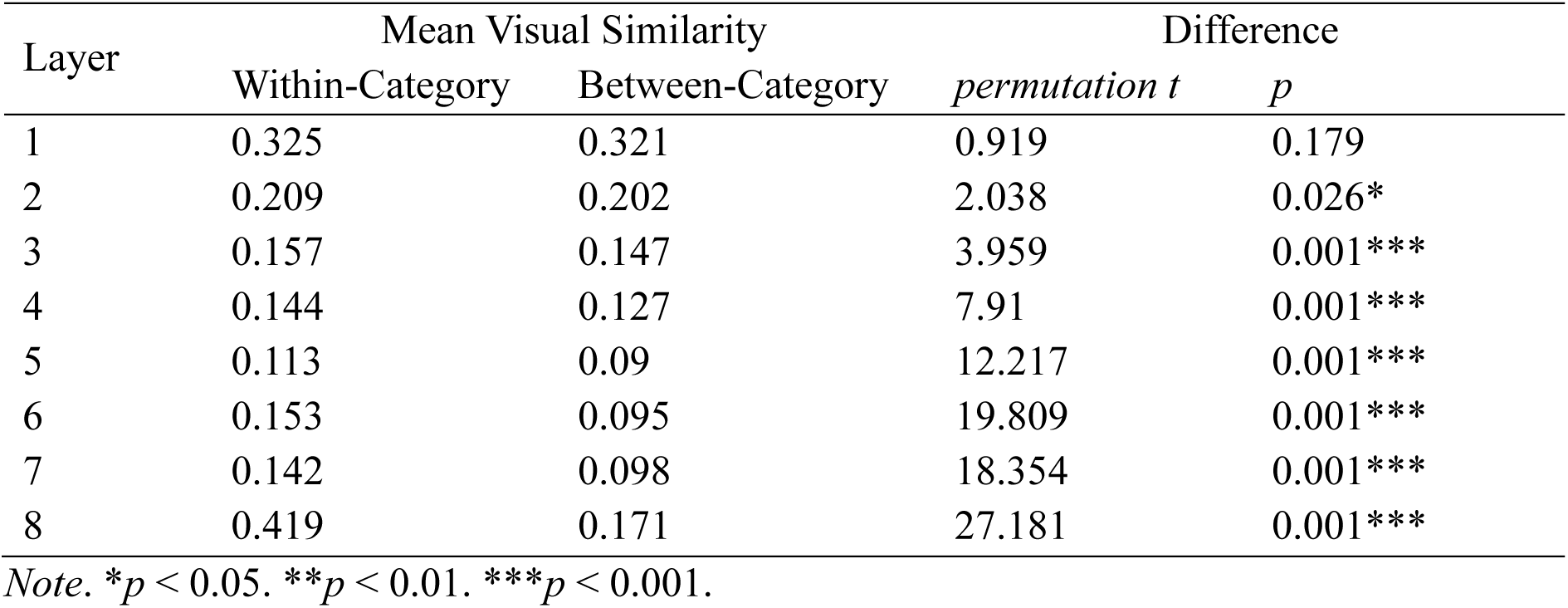
Mean within- and between-category visual similarity for each DNN layer and the difference in the mean similarity between within- and between-category pairs.

| Layer | Mean Visual Similarity |  | Difference |  |
| --- | --- | --- | --- | --- |
|  | Within-Category | Between-Category | permutation *t* | *p* |
| 1 | 0.325 | 0.321 | 0.919 | 0.179 |
| 2 | 0.209 | 0.202 | 2.038 | 0.026* |
| 3 | 0.157 | 0.147 | 3.959 | 0.001*** |
| 4 | 0.144 | 0.127 | 7.91 | 0.001*** |
| 5 | 0.113 | 0.09 | 12.217 | 0.001*** |
| 6 | 0.153 | 0.095 | 19.809 | 0.001*** |
| 7 | 0.142 | 0.098 | 18.354 | 0.001*** |
| 8 | 0.419 | 0.171 | 27.181 | 0.001*** |

*Note.* \* $p < 0.05$ . \*\* $p < 0.01$ . \*\*\* $p < 0.001$ .

#### Relating visual and semantic similarity to behavioural performance

To reveal unique relations between behavioural performance and visual/semantic similarity, we regressed out effects of one variable before estimating the effect of another variable, using two-step regression analyses. For both tasks, we first used the visual similarity of each layer to predict RTs under each set size condition separately, and then used categorical and/or semantic similarity to predict the residuals obtained in the first step. The same method and steps were used to test the relationship between visual similarity and behavioural performance. For each participant, each layer, and set size, all variables (including RT) were standardized using Z-score normalization. As a result, all the data were rescaled to have a mean of 0 and a standard deviation of 1.

## Results

### Analyses of mean ACC and RT

In the first analysis, we aimed to test how category influences memory search performance across different memory set sizes, replicating the category effects in previous studies (Drew & Wolfe, 2014; Shang et al., 2024), and to compare category and memory set size influences between LTM and STM search. Figure 3 shows the mean data for target-absent trials of the two tasks, and Table 2 shows the results of the three-way mixed ANOVAs with one between- (LTM/STM task) and two within-subject factors (within-/between-category condition; MSS 1/2/4/8). For both ACC and RT, all main effects and interactions were significant (Figure 3). Two-way ANOVAs for RT further revealed significant effects of MSS and category in both tasks (see Table 3). The results observed in the LTM task replicated the findings of a previous study (Shang et al., 2024) and showed that LTM and STM shared a similar pattern of set-size related RT increases. The analysis on ACC in the STM task mirrored these findings, while the effects on ACC in the LTM task were not significant. This was expected, as participants were close to ceiling in all LTM conditions (Figure 3).

**Figure 3.**
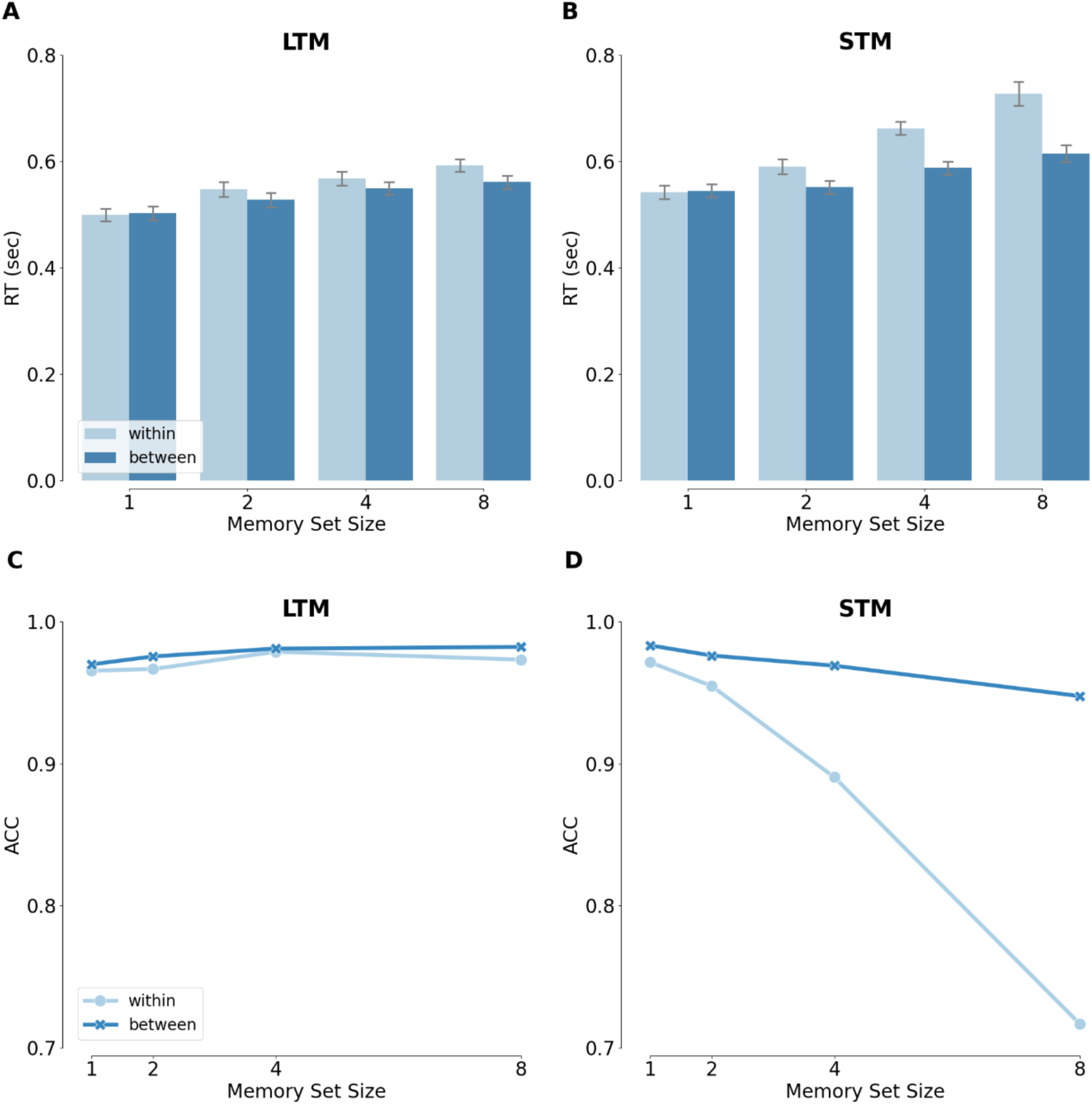
Mean performance (ACC and RT (in sec)) of target-absent trials, separately for LTM and STM tasks. *Note*. (A) and (B) show the mean RTs in LTM and STM tasks, respectively. (C) and (D) show the mean ACCs in LTM and STM tasks, respectively. Light blue indicates the within-category condition while dark blue indicates the between-category condition. For all figures, the error bars represent the standard error of the mean.

**Table 2.**
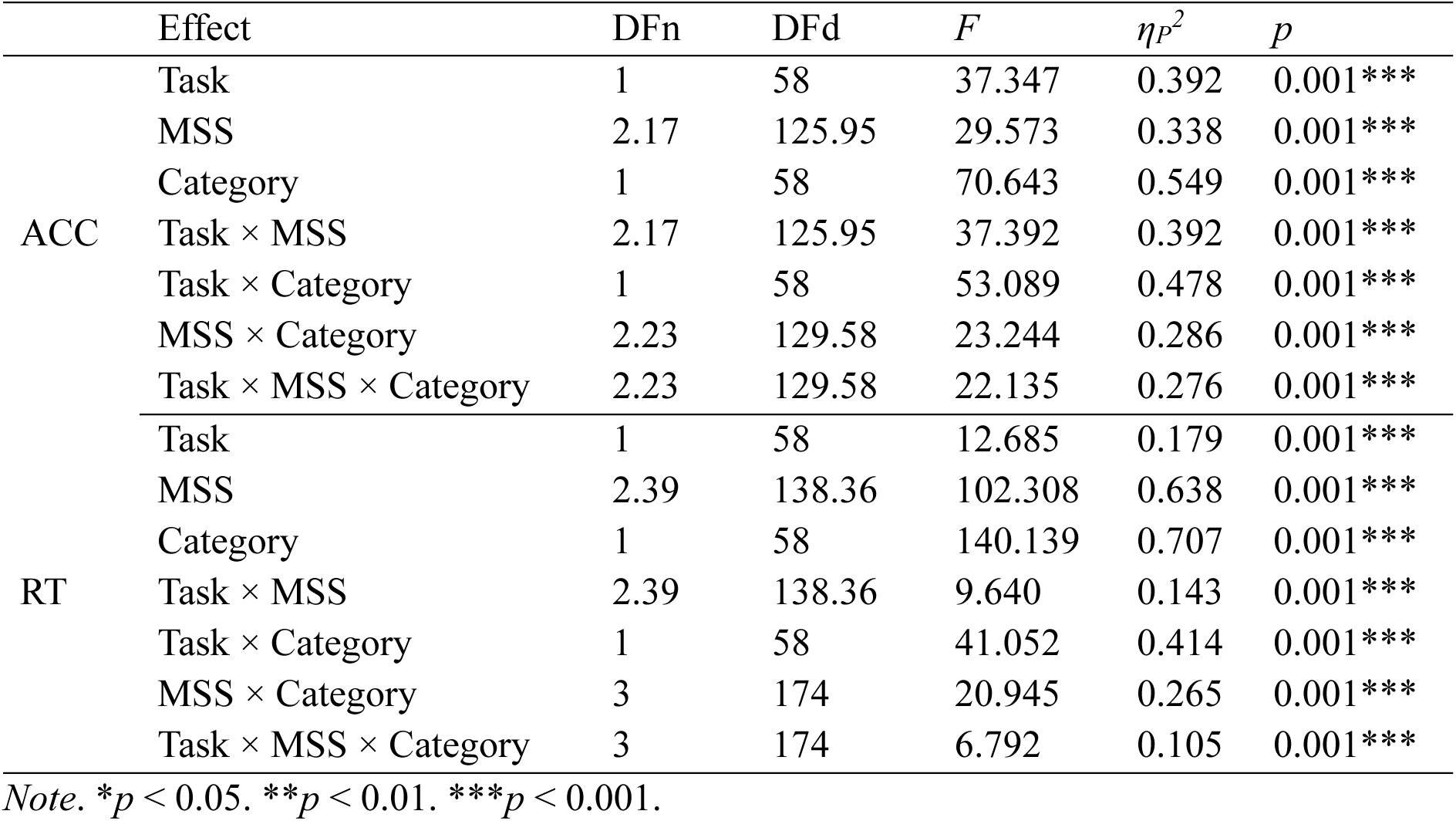
Three-way mixed ANOVA for ACC and RT (in sec).

**Table 3.**
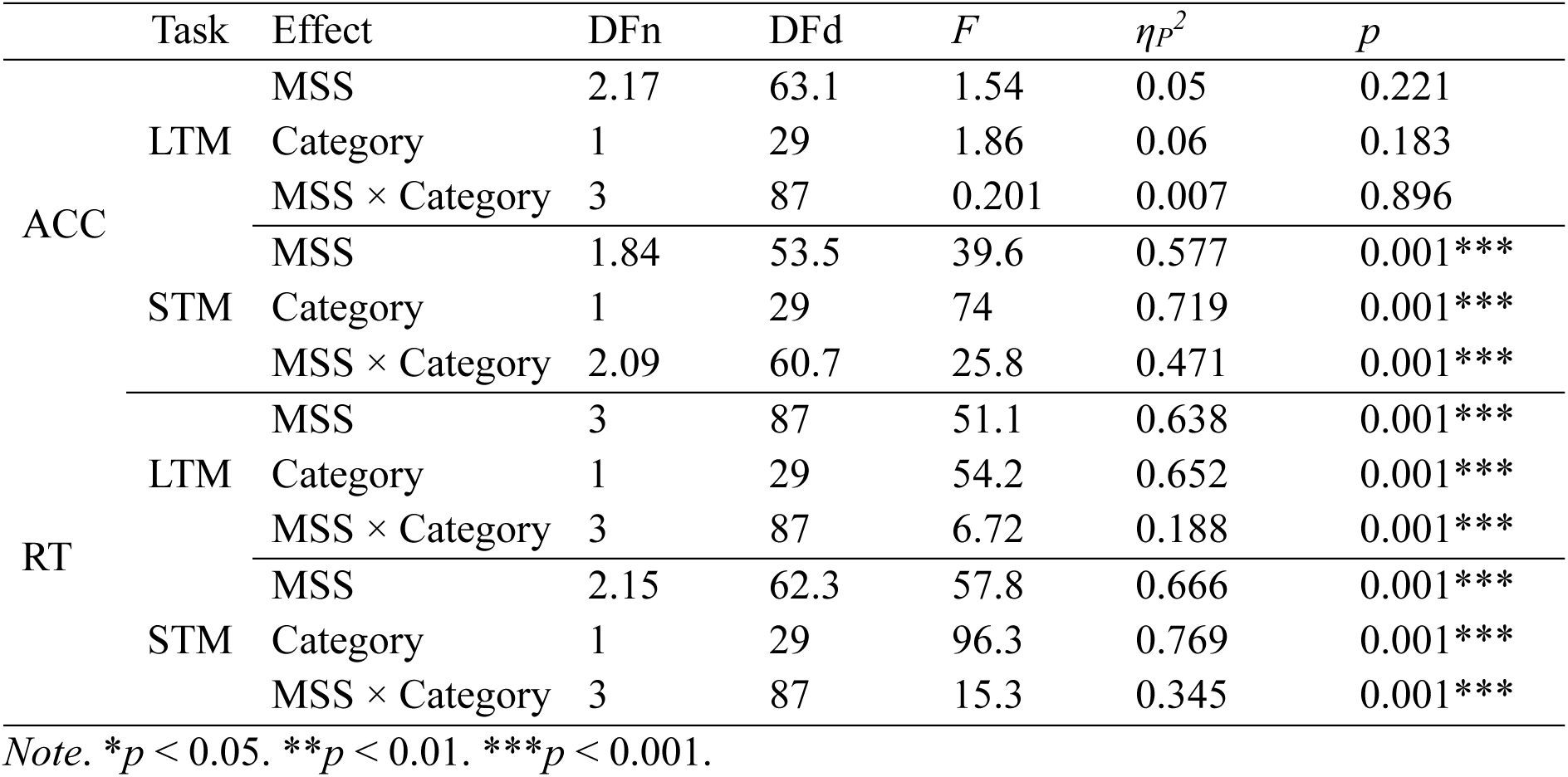
Two-way repeated ANOVA for ACC and RT (in sec), separately for LTM and STM tasks.

| | Task | Effect | DFn | DFd | *F* | $\eta_p^2$ | *p* |
| --- | --- | --- | --- | --- | --- | --- | --- |
| ACC | LTM | MSS | 2.17 | 63.1 | 1.54 | 0.05 | 0.221 |
|  |  | Category | 1 | 29 | 1.86 | 0.06 | 0.183 |
| | | MSS × Category | 3 | 87 | 0.201 | 0.007 | 0.896 |
|  | STM | MSS | 1.84 | 53.5 | 39.6 | 0.577 | 0.001*** |
|  |  | Category | 1 | 29 | 74 | 0.719 | 0.001*** |
| | | MSS × Category | 2.09 | 60.7 | 25.8 | 0.471 | 0.001*** |
| RT | LTM | MSS | 3 | 87 | 51.1 | 0.638 | 0.001*** |
|  |  | Category | 1 | 29 | 54.2 | 0.652 | 0.001*** |
| | | MSS × Category | 3 | 87 | 6.72 | 0.188 | 0.001*** |
|  | STM | MSS | 2.15 | 62.3 | 57.8 | 0.666 | 0.001*** |
|  |  | Category | 1 | 29 | 96.3 | 0.769 | 0.001*** |
| | | MSS × Category | 3 | 87 | 15.3 | 0.345 | 0.001*** |

*Note.* \* $p < 0.05$ . \*\* $p < 0.01$ . \*\*\* $p < 0.001$ .

To test whether RTs were log-linearly related to set size, following previous studies (e.g., Drew & Wolfe, 2014; Shang et al., 2024), the RTs from set size 1 to 4 were used to predict the performance on set size 8 (Figure 4A & B), For both tasks, the absolute error of the log-linear model was significantly smaller than the absolute error of the linear model, for both within-category, LTM: *t*_(29)_ = -3.895, *p* = 0.001, *d* = 0.711, STM: *t*_(29)_ = -4.176, *p* < 0.001, *d* = 0.762, and between-category conditions, LTM: *t*_(29)_ = -4.703, *p* < 0.001, *d* = 0.859, STM: *t*_(29)_ = -3.735, *p* = 0.001, *d* = 0.682.

**Figure 4.**
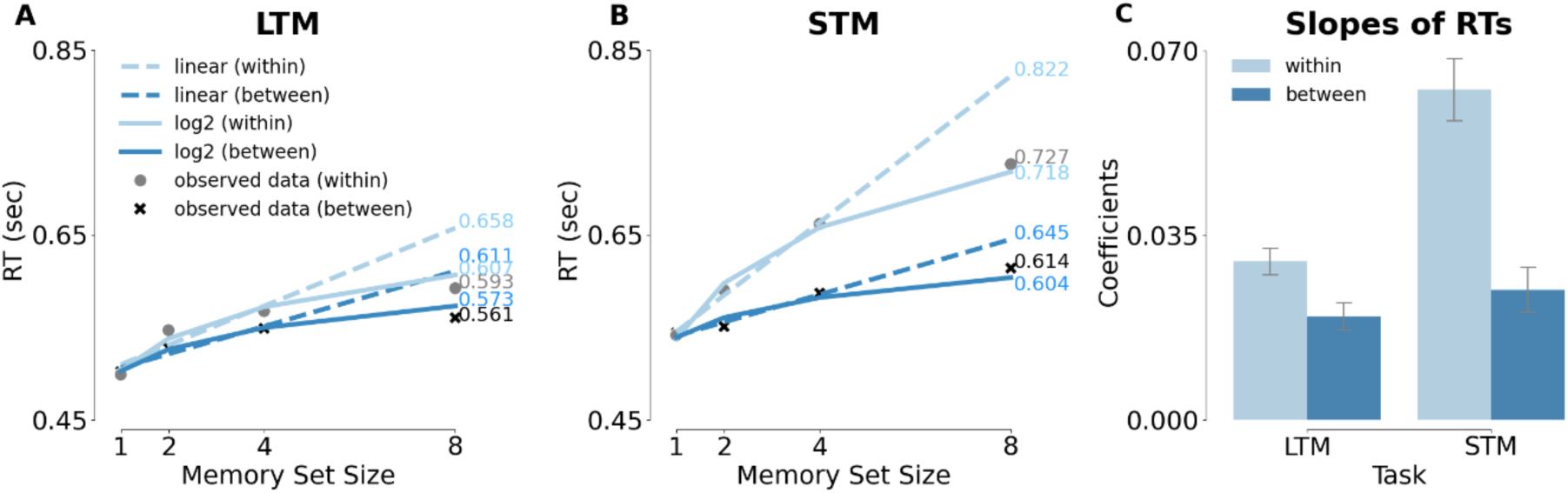
Linear vs log 2 model in target-absent trials. *Note*. (A) and (B) Linear vs log 2 model in target-absent trials of LTM and STM tasks. The light grey dots and dark grey crosses refer to the observed data in within-category and between-category conditions, respectively. The dashed lines represent linear regression models, while the solid lines represent log-linear models. For all the figures, the light blue always represents the within-category condition while the dark blue represents the between-category condition. (C) Log 2 slopes of the RTs in target-absent trials of LTM and STM tasks. The error bars represent the standard error of the mean.

Altogether, these results indicate that performance on the two tasks was similarly influenced by category and memory set size, although they differed in some aspects, as shown by two-way mixed ANOVAs on the log-linear slope coefficients. This analysis showed significant main effects of task and category condition, *F*_(1, 58)_ = 16.065, *p* < 0.001, *η^2^* = 0.217 and *F*_(1, 58)_ = 54.494, *p* < 0.001, *η^2^* = 0. 484. There was also a significant interaction between task and category condition, *F*_(1, 58)_ = 17.672, *p* < 0.001, *η^2^* = 0. 234, reflecting a stronger category effect in STM than in LTM (Figure 4C). Importantly, however, the simple main effects of category were significant in both tasks, LTM: *F*_(1, 29)_ = 18.1, *p* < 0.001, *η^2^*= 0. 384 and STM: *F*_(1, 29)_ = 39, *p* < 0.001, *η^2^*= 0. 574, with steeper slopes (less efficient memory search) for within-category distractors (Figure 4).

### Analyses of visual similarity

#### Basic visual similarity

One of the main aims of the current study was to investigate whether category can explain the performance on memory recognition beyond visual similarity. To this end, we first tested if trial-wise RT variability in memory search could be explained by target-distractor visual similarity. The sklearn and statsmodels.api packages in Python were employed to perform GLM. The z-transformed visual similarity established using AlexNet (see Methods) was employed to predict the z-transformed reaction times of old/new recognition tasks using regression analysis, separately for each set size. To assess the statistical significance of the beta values obtained from the GLM, we performed a one-sample t-test at each AlexNet layer across participants and applied a cluster-based permutation test to correct for multiple comparisons over layers. Given that the eight AlexNet layers are sequentially ordered and reflect the hierarchical organization of the neural network, adjacency was defined such that layer 1 is considered adjacent to layer 2, layer 2 adjacent to layer 3, and so on. Layers with p-values < 0.05 were grouped together to form clusters. To generate a null distribution, the signs of the beta values were randomly flipped, and the t-tests were recomputed across 1000 permutations. For each permutation, the largest cluster-level statistic was retained, forming a distribution of maximum cluster statistics under the null hypothesis. The cluster statistics obtained from the observed data were compared against the null distribution generated through permutations to calculate p-values. Clusters with p-values below the threshold (i.e., 0.05) were considered statistically significant. The permutation_cluster_1samp_test package based on MNE in Python was employed to perform the cluster-based permutation test. The results showed that visual similarity could significantly predict RT in both tasks (cluster-based *p* < 0.05). For both tasks, visual similarity in the late AlexNet layers best predicted RT, with a peak in the 6^th^ layer (Figure 5).

**Figure 5.**
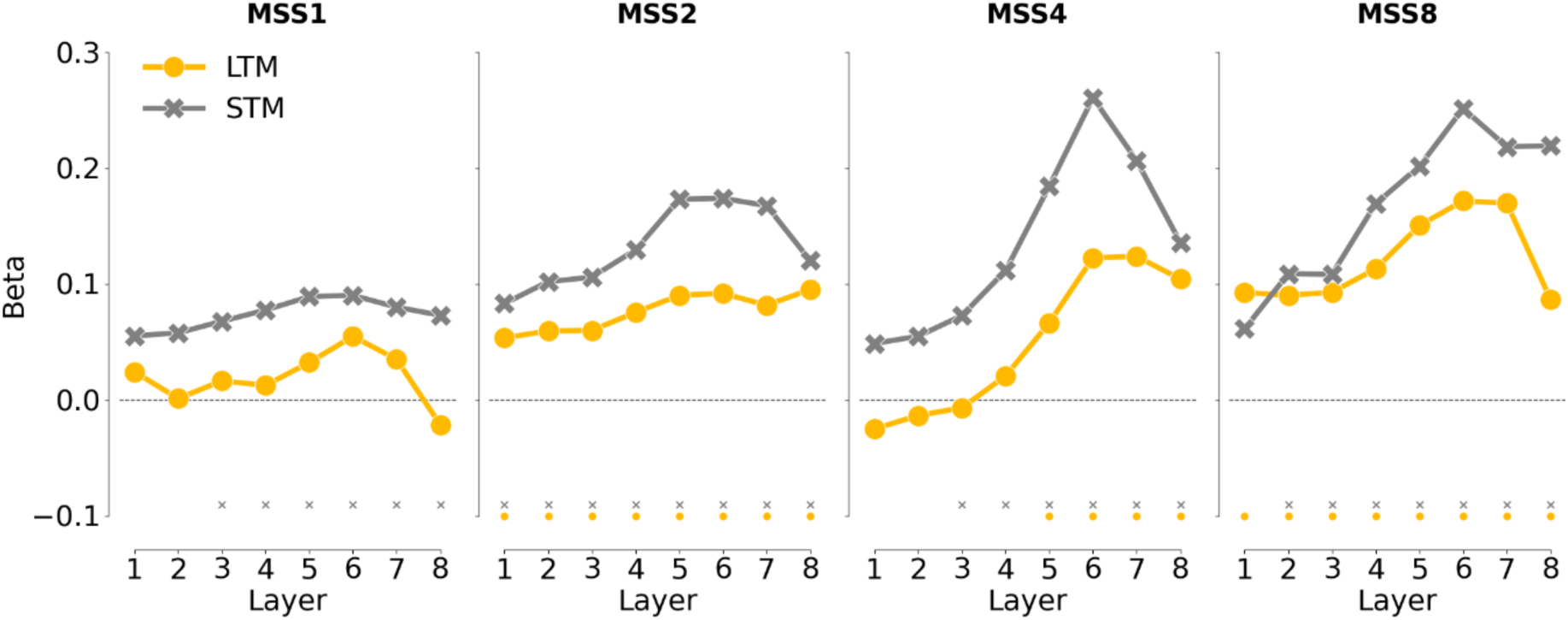
Results of regression analyses for visual target-distractor similarity. *Note*. Visual similarity (derived from the 8 layers of AlexNet) was used to predict trial-wise RTs for the two tasks and the four memory set sizes (MSS) separately. The orange lines represent results of the LTM task while the grey lines represent results of the STM task. **Asterisks indicate significance based on the cluster-based permutation test (\**p* < 0.05).**

Considering that Layer 6 best predicted RT across conditions, we followed up on this layer, comparing the visual similarity across the four set sizes and the two tasks. A two-way mixed ANOVA (LTM/STM task × MSS 1/2/4/8) showed significant main effects of task, *F*_(1, 58)_ = 14.085, *p* < 0.001, *ηP*^2^ = 0.195, and MSS, *F*_(2.91, 168.7)_ = 9.419, *p* < 0.001, *ηP*^2^ = 0.140, but no significant interaction, *F*_(2.91, 168.7)_ = 1.057, *p* = 0.368, *ηP*^2^ = 0.018. These results indicate that Layer-6 visual similarity more strongly predicted RTs in the STM than the LTM task (Figure 6A). Furthermore, as memory set size increased, the contribution of visual similarity gradually became larger. Finally, the influence of visual similarity in LTM and STM tasks followed a similar set size pattern.

**Figure 6.**
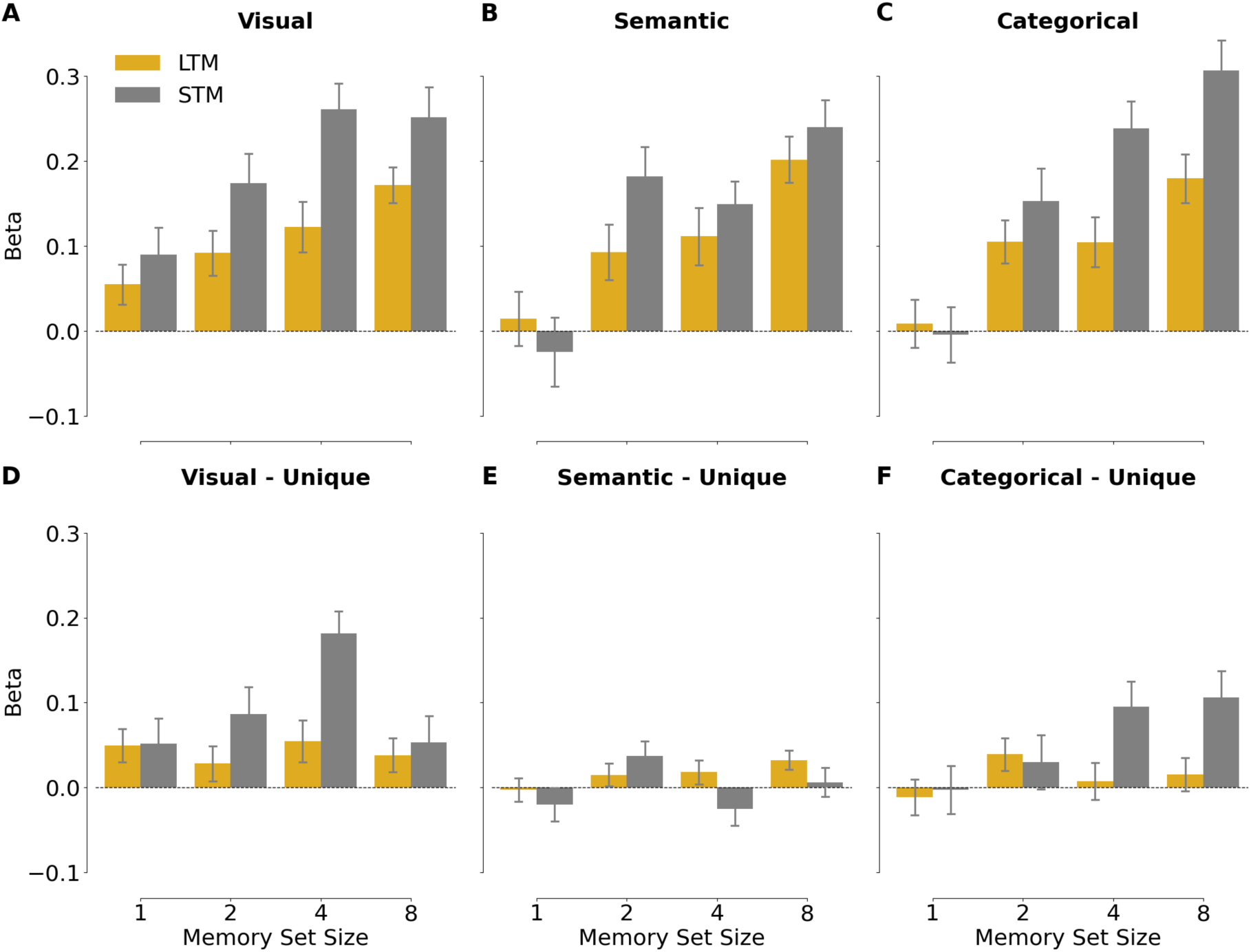
Results of regression analyses for Layer-6 target-distractor similarity. *Note*. Visual - Unique = visual similarity after regressing out categorical and semantic similarity; Semantic - Unique = semantic similarity after regressing out visual and categorical similarity. Categorical - Unique = categorical similarity after regressing out visual and semantic similarity. The orange bars represent results of the LTM task while the grey bars represent results of the STM task.

#### Visual similarity after accounting for categorical and semantic similarity

In order to reveal the unique contribution of Layer-6 visual similarity to trial-wise RT variability, semantic similarity was first used to predict RTs separately in the within- and between-category conditions. Then, visual similarity was used to predict the residuals obtained in the first step, separately for the two categories. Finally, the results were averaged across the two categories.

A two-way mixed ANOVA for the visual similarity beta values (LTM/STM task × MSS 1/2/4/8) showed main effects of task, *F*_(1, 58)_ = 7.711, *p* = 0.007, *ηP*^2^ = 0.117, and MSS, *F*_(2.9, 168.3)_ = 3.355, *p* = 0.022, *ηP*^2^ = 0.055, while the interaction did not reach the significance level, *F*_(2.9, 168.3)_ = 2.331, *p* = 0.078, *ηP*^2^ = 0.039. These findings again showed a stronger influence of visual similarity in predicting STM than LTM performance (Figure 6D).

### Analyses of semantic similarity

#### Basic semantic similarity

In the second series of analyses, we asked whether the semantic similarity of target and distractor objects predicted trial-wise RT variability. A two-way mixed ANOVA showed a significant main effect of MSS, *F*_(2.74, 158.9)_ = 15.506, *p* < 0.001, *ηP*^2^ = 0.211. The main effect of task, *F*_(1, 58)_ = 2.18, *p* = 0.145, *ηP*^2^ = 0.036, and the interaction were not significant, *F*_(2.74, 158.9)_ = 1.249, *p* = 0.294, *ηP*^2^ = 0.021. This result indicates that the influence of semantic similarity increases with set size but does so similarly for STM and LTM (Figure 6B).

#### Semantic similarity after accounting for visual and categorical similarity

In order to reveal the unique contribution of semantic similarity to trial-wise RT variability, visual similarity (Layer-6) was first used to predict RTs separately in the within- and between-category conditions. Then, semantic similarity was used to predict the residuals obtained in the first step, separately for the two categories. Finally, the results were averaged across the two categories.

A two-way mixed ANOVAs showed no significant main effects or interaction, task: *F*_(1, 58)_ = 1.954, *p* = 0.168, *ηP*^2^ = 0.033; MSS: *F*_(2.89, 167.3)_ = 2.541, *p* = 0.06, *ηP*^2^ = 0.042, interaction: *F*_(2.89, 167.3)_ = 1.511, *p* = 0.215, *ηP*^2^ = 0.025 (Figure 6E).

### Analyses of categorical similarity

#### Basic categorical similarity

In the final section, we turn to the question of whether category (as a binary variable, animate vs inanimate) explains the performance on memory recognition beyond visual and semantic similarity. When not regressing out visual similarity, categorical similarity was a good predictor of memory performance (Figure 6C). An ANOVA showed that there were significant differences in the category beta estimates between the two tasks, *F*_(1, 58)_ = 14.686, *p* < 0.001, *ηP*^2^ = 0.202. The main effect of MSS was also significant, *F*_(2.91, 168.6)_ = 19.174, *p* < 0.001, *ηP*^2^ = 0.248. However, the interaction was not significant, *F*_(2.91, 168.6)_ = 2.289, *p* = 0.082, *ηP*^2^ = 0.038, suggesting a similar MSS-related increase in the two tasks.

#### Categorical similarity after regressing out visual and semantic similarity from RT

We repeated these analyses after regression out visual and semantic similarity. In this case, only the main effect of task was significant, *F*_(1, 58)_ = 7.113, *p* = 0.01, *ηP*^2^ = 0.109; MSS: *F*_(2.69, 155.9)_ = 2.518, *p* = 0.067, *ηP*^2^ = 0.042, and interaction: *F*_(2.69, 155.9)_ = 1.908, *p* = 0.137, *ηP*^2^ = 0.032. These results indicate that compared with LTM search, in which the contribution of superordinate category was relatively weak, STM search depended more on the binary categorical distinction (i.e., animacy), especially in large set size (Figure 6F).

#### Comparison of the contributions of visual, semantic, and categorical similarity between LTM and STM tasks

The previous sections showed no interactions between task and set size, indicating that the influence of visual, semantic, and categorical similarity similarly changed across set size for the two tasks. Therefore, in a final analysis, we averaged across the set size conditions to compare the unique influence of visual (Layer-6), semantic, and categorical similarity between the tasks. Figure 7 summarizes the contributions to performance in the two memory tasks of unique visual (regressing out categorical and semantic similarity), unique semantic (regressing out categorical and visual similarity), and unique categorical (regressing out visual and semantic similarity) similarity. Because betas cannot be directly compared across dimensions, two-sample t-tests were employed to compared the unique similarity influences across the two tasks. Results showed stronger influences on STM than LTM performance of visual similarity, *t*_(49.791)_ = -2.777, *p* = 0.008, *d* = -0.192, and categorical similarity, *t*_(47.876)_ = - 2.667, *p* = 0.010, *d* = -0.165 (Figure 7A and C). By contrast, the influence of semantic similarity was significant only for LTM (*t*_(29)_ = -2.80, *p* = 0.009) and was numerically larger for LTM than STM (Figure 7B), although this difference was not significant, *t*_(45.343)_ = 1.398, *p* = 0.169, *d* = 0.870.

**Figure 7.**
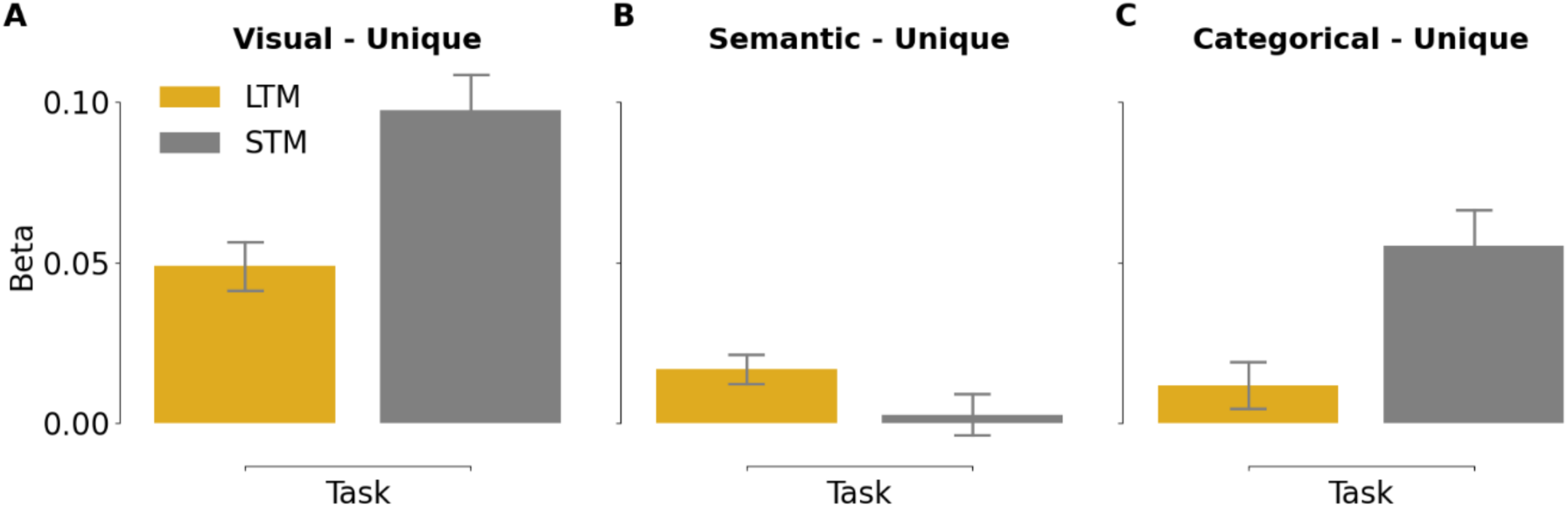
Comparison of the mean (averaged across set size) unique contributions of visual, categorical, and semantic similarity between the two tasks. *Note*. Visual - Unique = visual similarity after regressing out categorical and semantic similarity; Semantic - Unique = semantic similarity after regressing out visual and categorical similarity. Categorical - Unique = categorical similarity after regressing out visual and semantic similarity. The orange bars represent results of the LTM task while the grey bars represent results of the STM task.

#### General discussion

In this study, we investigated the roles of visual, semantic, and categorical similarity in short- and long-term memory search. While these three factors are correlated, they are clearly not identical. Indeed, our results show that all factors predicted trial-wise variability in reaction times, even after the other factors were regressed out or controlled for. The results answer the three questions outlined in the Introduction: First, results showed that the category effect in long-term memory search, as reported in previous studies (e.g., Gronau et al., 2024; Shang et al., 2024; Williams et al., 2024; Cunningham & Wolfe, 2012, 2014; Drew & Wolfe, 2014), remained significant even when visual and semantic similarity were regressed out. Second, we found that visual and semantic similarity also independently contributed to long-term memory search performance. Finally, we found that LTM and STM differed in the contributions of visual, semantic, and categorical similarities: STM was more strongly influenced by visual and categorical similarity than LTM was, while this was not the case for semantic similarity (Figure 7). We will discuss each of these findings below.

#### Category effects in LTM search

Previous studies observed significant categorical influences on memory search performance that increased with increasing set size (Gronau et al., 2024; Shang et al., 2024; Williams et al., 2024; Cunningham & Wolfe, 2012, 2014; Drew & Wolfe, 2014), which we replicated here. However, none of these studies investigated what exactly contributed to this category effect, specifically, whether the category effects could be explained by object attributes (e.g., visual or semantic target-distractor similarity) other than the binary categorical target-distractor similarity. Therefore, in the current study, we used neural network and distributional semantic models to establish visual similarity and semantic similarity, respectively, in order to measure (and regress out) their influence on memory search performance. We observed that when the influences from visual and semantic similarity were regressed out, the superordinate category of the distractor still predicted long-term memory search performance, confirming the distinct contribution of task-related category membership.

Using category membership is an efficient strategy to quickly reject distractors when all targets are drawn from the same category. This could be achieved by directing attention to visual features that are diagnostic of the target category (e.g., curved shapes when all memory items are animals), such that objects from other categories can be filtered out before the memory search stage (Shang et al., 2024). This account predicts that reaction times scale with the degree to which an object matches the attentional template (Duncan & Humphreys, 1989). However, the features included in the template are not known. One possibility is that the template consists of the visual features that are common to the items in the memory set. In that case, rejecting distractor probe items that are visually relatively similar to the items in the memory set would be relatively slow. In the current experiment, this variability is captured by our model of the average visual similarity of the probe item to the items in the memory set, which was a good predictor of trial-wise RT variability (Figure 7). Indeed, regressing out high-level visual similarity reduced the category effect.

Interestingly, we observed a significant residual category effect when regressing out the contribution of visual similarity. This raises the possibility that participants formed a more abstract (or generic) categorical template, based on experience with that category (e.g., animals) across a lifetime. When forming an attentional template, it may be easier for participants to activate an overlearned categorical representation than to retrieve the shared features of the items in the memory set, which vary from trial to trial (in STM) or from block to block (in LTM). In line with this interpretation, previous studies have provided evidence that attention can be directed at the level of object category for highly familiar categories (Peelen et al., 2014) including animals (Evans & Treisman, 2005; Stein & Peelen, 2017).

#### Visual and semantic similarity independently predict LTM search

In addition to category, memory search performance was also significantly and independently predicted by visual similarity across AlexNet layers, with a peak at the 6^th^ layer (Figure 5). The 6^th^ layer acts as a bridge between convolutional feature maps and classification. It contains a high-dimensional representation of high-level visual features that supports object classification. This representation may be similar to the transformation from perceptual to categorical representations in higher-order visual cortex areas. As discussed above, the influence of visual similarity could reflect the partial match of distractor items to the attentional template, with visually highly dissimilar distractors being rejected already before the memory search stage. Alternatively, it could reflect interference during the memory search itself, with visually similar distractors being more effective lures than visually dissimilar distractors.

When the influences of visual and categorical similarity were regressed out, semantic similarity also still predicted LTM search performance (Figure 7B). This result is in line with previous work that emphasized the role of conceptual distinctiveness in memory recognition (Konkle et al., 2010b). Support from conceptual structure was shown to contribute to explaining the remarkable performance on visual memory tasks involving thousands of pictures (Konkle et al., 2010b). Our finding that visual and semantic similarity both contributed to memory task performance is also in line with other recent studies. For example, participants were shown to depend on both visual and semantic information to classify pictures, such that when pictures had similar semantics, participants took longer to classify them (Lotto et al., 1999). Furthermore, compared with visual information, semantic information plays a greater role in memory tasks (Shoham et al., 2024). Shoham et al. (2024) also reported that when participants recalled images from memory, they depended more on semantic information to reconstruct the visual features of the images. However, while these studies emphasized the important role of semantic information in memory, they also highlighted that through mental representations, visual and semantic information interactively influence perception and memory tasks.

#### Comparing LTM and STM

Before turning to the contributions of categorical, semantic, and visual similarity, it is worth nothing the similar log-linear set-size-RT increase in STM and LTM (Figure 4). While previous studies also found such a log-linear set-size-RT increase, this was interpreted as an effect of the study-test lag resulting from the serial presentation of stimuli (Nosofsky, Cao, et al., 2014; Nosofsky, Cox, et al., 2014). In our study, with parallel presentation in the STM task, and analyzing responses to unique distractor stimuli in both tasks, study-test lag differences could not account for the log-linear increase. Furthermore, the similarity contributions to LTM and STM followed surprisingly similar patterns across set size. These findings suggest partly shared mechanisms of search in STM and LTM (see also Schurgin, 2018).

Averaged across set size, unique semantic similarity predicted LTM but not STM performance (Figure 7B), while, conversely, the influences of visual similarity and categorical similarity were greater for STM than LTM (Figure 7A and C). We found that semantic similarity strongly and reliably predicted STM performance when it was the only predictor in the model (Figure 6B). Further analyses, however, revealed that the effect of semantic similarity could be fully accounted for by a combination of visual and categorical similarity; when regressing out visual and categorical similarity, semantic similarity no longer explained variance in RT data in the STM task (Figure 6E). This finding highlights the importance of accounting for correlated stimulus dimensions, as done here using regression analysis. To further reveal their separate and interactive influence on memory performance, future studies could use stimulus sets in which visual, semantic, and categorical similarity are experimentally manipulated (e.g., Yeh & Peelen, 2022).

It should be noted that the lack of a unique semantic similarity contribution to STM performance does not imply that semantic information does not contribute to STM at all. For example, previous studies demonstrated that STM performance is better for meaningful than meaningless stimuli (Sahar et al., 2020; 2024; Shoval and Makovski, 2022), which could also influence the upper limit of working memory capacity (Asp et al., 2021). By contrast to these studies, our study focused on the influence of semantic target-distractor similarity on trial-wise RT, showing that when the influence of visual and categorical similarity are regressed out, semantically similar and dissimilar distractor probes are rejected equally fast. Finally, the finding that the contribution of visual similarity was more important for STM than for LTM reinforces the idea that STM uses a more perceptual code than LTM to represent objects (for reviews, see: Luck, 2008; Sreenivasan et al., 2014).

#### Special cases

Set size 1 is a special case in our experiment, as for this set size, the task is effectively a single-target detection task rather than a memory search task. Participants can maintain a precise top-down attentional template for one item (Olivers et al., 2011) such that all distractor objects can be quickly rejected, irrespective of their semantic or categorical similarity to the target item. Accordingly, only visual similarity explained trial-wise RT variability (Figure 5). Nevertheless, when there is competition (e.g., when presenting multiple probe items), semantic and categorical similarity could still influence visual orienting later in time (De Groot et al., 2016; Yeh & Peelen, 2022).

Set size 8 is a special case for STM because the set size exceeds the capacity of visual STM (∼4 items). In this case, categorical similarity had a particularly strong influence on RT (Figure 6 C&F). This may be because participants could only memorize the animacy of the target objects, and then rejected the between-category distractors primarily based on their category. This would also lead to high error rates for within-category distractors compared with between-category distractors, as confirmed by the accuracy data, *t*_(29)_ = 7.602, *p* < 0.001, *d* = 1.388, which was not observed for set size 8 of the LTM task, *t*_(29)_ = 1.137, *p* = 0.265, *d* = 0.208. This suggests that categorical strategies compress visual information (Cowan, 2001; Olsson & Poom, 2005). For example, Olsson and Poom (2005) demonstrated that when the stimuli belonged to different categories (e.g., discrete colour/shape), participants could remember nearly 3 stimuli, while when all the objects belonged to a single category, only one stimulus could be remembered because all the information was compressed into one category which impeded the recognition of target objects. This indicates that STM requires the support of categorical structures in LTM. Participants in the STM task may remember categories rather than objects. Thus, participants can quickly reject the distractor objects from different categories through categorial similarity.

## Conclusion

In summary, the current study shows that memory search performance is determined by a combination of visual, semantic, and categorical target-distractor similarity. For both LTM and STM tasks, visual, semantic, and categorical similarity each contributed to predicting memory search performance, following similar patterns across set size in both tasks. However, visual and categorical influences (but not semantic influences) were stronger in STM than LTM search. These results clarify the nature of the representations used in STM and LTM search.

