## Supplementary Material for "Unpacking similarity effects in visual memory search: categorical, semantic, and visual contributions"

### Results based on maximum visual similarity

The regression results based on maximum visual similarity (see Figure S1) were almost identical to the findings based on mean visual similarity. Similar to the analysis using mean similarity, maximum similarity at Layer 6 also best predicted RT across conditions. Accordingly, we followed up on this layer, comparing the visual similarity across the four set sizes and the two tasks. A two-way mixed ANOVA (LTM/STM task × MSS 1/2/4/8) showed significant main effects of task, $F_{(1, 58)} = 13.039$, $p < 0.001$, $\eta_P^2 = 0.184$, and MSS, $F_{(2.91, 168.3)} = 8.995$, $p < 0.001$, $\eta_P^2 = 0.134$, but no significant interaction, $F_{(2.91, 168.3)} = 0.536$, $p = 0.652$, $\eta_P^2 = 0.009$.

### Figure S1

*Results of regression analyses for visual target-distractor similarity based on maximum similarity*

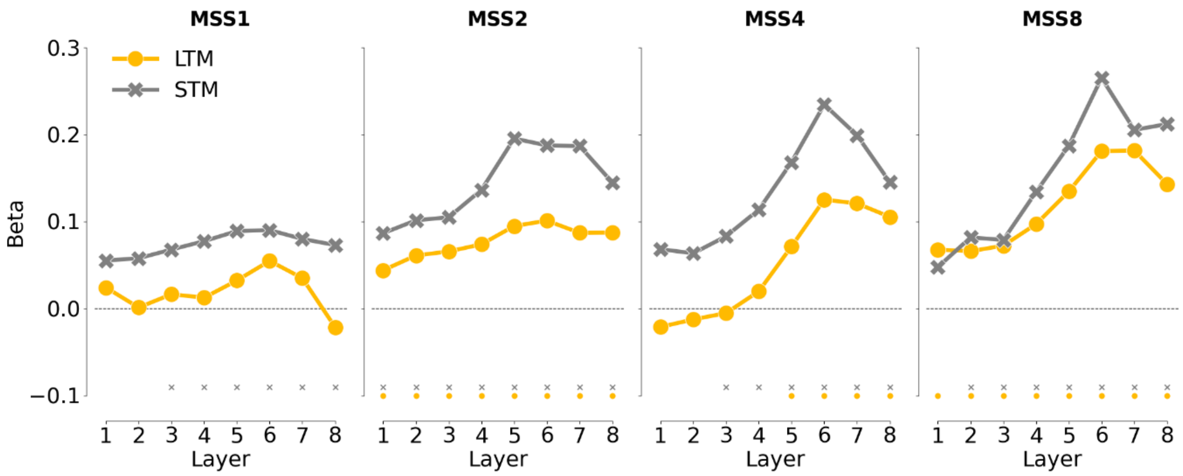

*Note*. Maximum visual similarity (derived from the 8 layers of AlexNet) was used to predict trial-wise RTs for the two tasks and the four memory set sizes (MSS) separately. The orange lines represent results of the LTM task while the grey lines represent results of the STM task. Asterisks indicate significance based on the cluster-based permutation test (*$p < 0.05$).

**Figure S2**

*Results of regression analyses for Layer-6 target-distractor similarity based on maximum similarity*

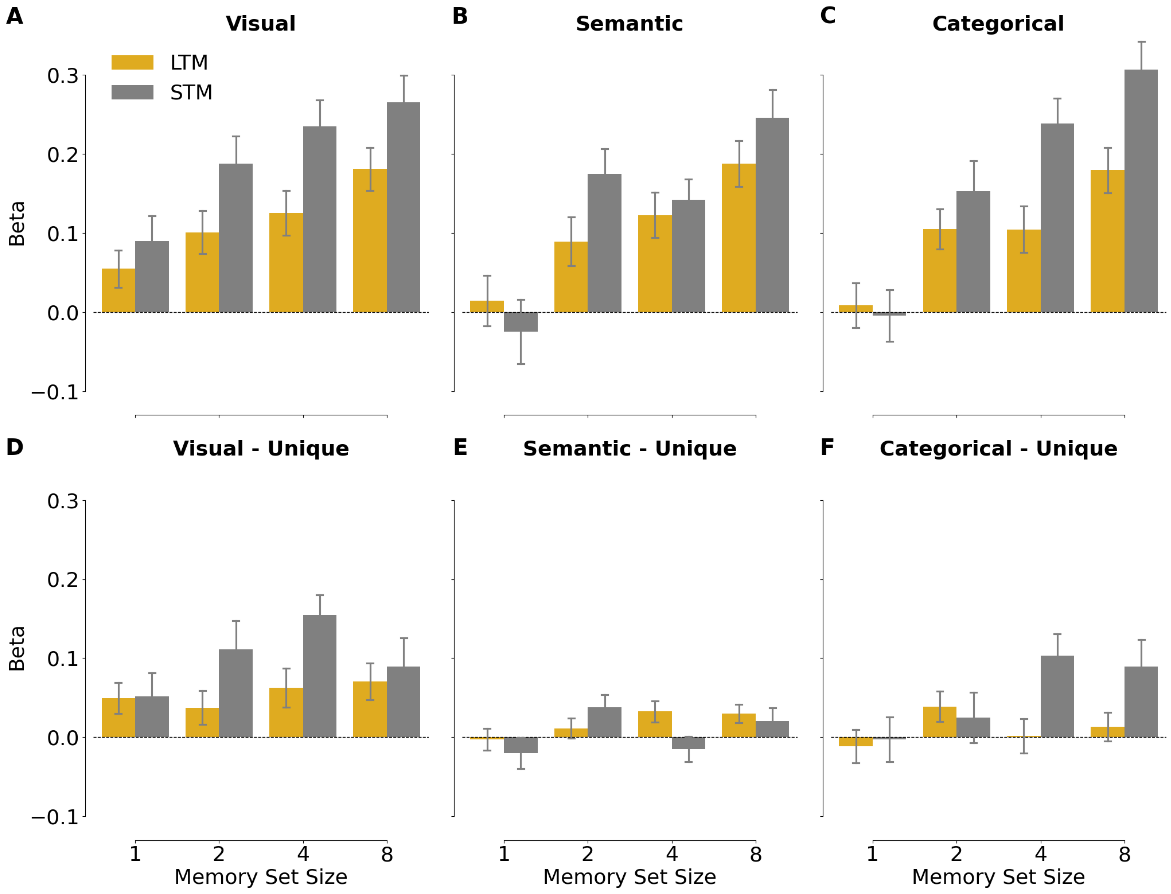


*Note*. Visual - Unique = visual similarity after regressing out categorical and semantic similarity; Semantic - Unique = semantic similarity after regressing out visual and categorical similarity. Categorical - Unique = categorical similarity after regressing out visual and semantic similarity. The orange bars represent results of the LTM task while the grey bars represent results of the STM task.

***Visual similarity after accounting for categorical and semantic similarity***

A two-way mixed ANOVA for the beta values at the 6th layer (LTM/STM task × MSS 1/2/4/8) showed a main effect of task only, $F_{(1, 58)} = 5.763$, $p = 0.02$, $\eta_P^2 = 0.09$, while MSS, $F_{(2.96, 171.7)} = 1.448$, $p = 0.231$, $\eta_P^2 = 0.024$, and the interaction, $F_{(2.96, 171.7)} = 1.181$, $p = 0.318$, $\eta_P^2 = 0.02$, did not reach the significance level. These findings again showed a stronger influence of visual similarity in predicting STM than LTM performance (Figure 6D).

**Analyses of semantic similarity**

***Basic semantic similarity***

A two-way mixed ANOVA showed a significant main effect of MSS, $F_{(2.64, 152.9)} = 15.336$, $p < 0.001$, $\eta_P^2 = 0.209$. The main effect of task, $F_{(1, 58)} = 2.294$, $p = 0.135$, $\eta_P^2 = 0.038$, and the interaction were not significant, $F_{(2.64, 152.9)} = 1.320$, $p = 0.271$, $\eta_P^2 = 0.022$.

***Semantic similarity after accounting for visual and categorical similarity***

A two-way mixed ANOVAs showed a main effect of MSS, $F_{(2.84, 164.5)} = 2.742$, $p = 0.048$, $\eta_P^2 = 0.045$. The main effect of task, $F_{(1, 58)} = 1.116$, $p = 0.295$, $\eta_P^2 = 0.019$, and the interaction, $F_{(2.84, 164.5)} = 2.130$, $p = 0102$, $\eta_P^2 = 0.035$, were not significant.

**Analyses of categorical similarity**

***Basic categorical similarity***

The results of maximum categorical similarity were identical to the results of mean similarity.

***Categorical similarity after regressing out visual and semantic similarity from RT***

The main effect of task was significant, $F_{(1, 58)} = 6.652$, $p = 0.012$, $\eta_P^2 = 0.103$; the other effects were not significant MSS: $F_{(2.85, 165)} = 2.198$, $p = 0.094$, $\eta_P^2 = 0.037$, and interaction: $F_{(2.85, 165)} = 2.123$, $p = 0.103$, $\eta_P^2 = 0.035$.

**Comparison of the contributions of visual, semantic, and categorical similarity between LTM and STM tasks**

Similar to the analyses using mean similarity, also for maximum similarity, results showed stronger influences on STM than LTM performance of visual similarity, $t_{(50)} = -2.401$, $p = 0.02$, $d = -0.099$, and categorical similarity, $t_{(45.576)} = -2.579$, $p = 0.013$, $d = -0.143$ (Figure S3A and C). By contrast, the influence of semantic similarity was numerically larger for LTM than STM (Figure S3B), although this difference was not significant, $t_{(46.356)} = 1.056$, $p = 0.296$, $d = 0.78$.

**Figure S3**

*Comparison of the mean (averaged across set size) unique contributions of maximum visual, categorical, and semantic similarity between the two tasks*

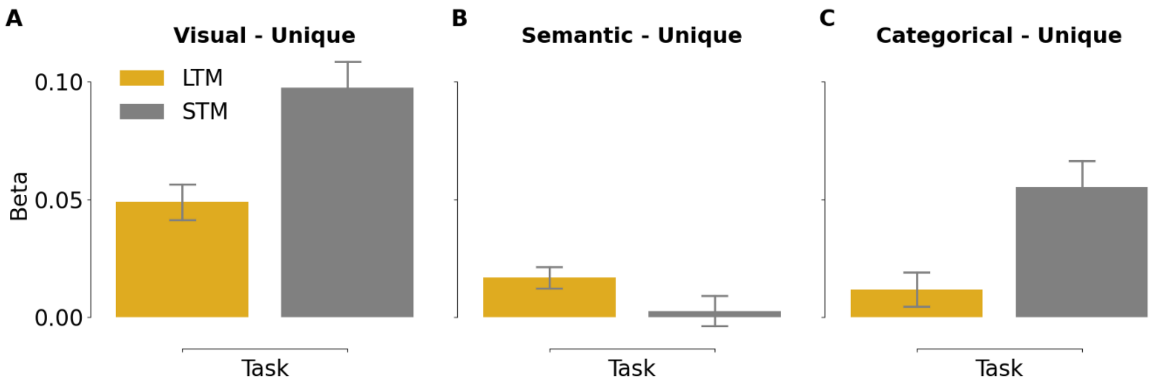


*Note*. Visual - Unique = maximum visual similarity after regressing out categorical and maximum semantic similarity; Semantic - Unique = maximum semantic similarity after regressing out maximum visual and categorical similarity. Categorical - Unique = categorical similarity after regressing out maximum visual and semantic similarity. The orange bars represent results of the LTM task while the grey bars represent results of the STM task.
